# Fecal Microbiota Transplantation Decreases Amyloid Load and Improves Cognition in Alzheimer’s

**DOI:** 10.1101/687376

**Authors:** Shalini Elangovan, Thomas J Borody, R M Damian Holsinger

## Abstract

The efficacy of fecal microbiota transplantation (FMT) in Alzheimer’s disease has yet to be investigated. Here, we show that FMT is capable of providing neuroprotective effects in two groups of treated 5xFAD Alzheimer’s mice, old transgenic (Tg) mice fed fecal slurry from healthy, wild-type donors of similar age (Old Tg-FO) and old mice fed fecal slurry from younger healthy, wild-type donors (Old Tg-FY). Improved spatial and recognition memory in Old Tg-FY and enhanced recognition memory in Old Tg-FO were observed when compared to Old Tg-Control mice given saline. Crucially, there was significant decreases in cortical Aβ loading in all treated mice, demonstrating the therapeutic effects of FMT in improving cognition and reducing amyloid pathology in AD brains.

**One Sentence Summary:** Fecal microbial transplants reduce amyloid pathology and improve cognition in Alzheimer’s mice.

## Main Text

Alzheimer’s disease (AD) is a progressive neurodegenerative condition resulting in cognitive, behavioral and functional decline with a predementia phase termed mild cognitive impairment (MCI) (*1, 2*). The pathological hallmarks of AD include extracellular deposition of amyloid-β (Aβ) peptide and intracellular accumulation of the hyperphosphorelated microtubule-associated protein tau into neurofibrillary tangles (NFT). Evidence suggests that these two pathological deposits are potentially involved in feedback loops where each causes further deposition of the other (*3, 4*) leading to synaptic degeneration (*5, 6*). Although genetic risk factors have been well-established in early-onset AD (*6–9*), various other causal mechanisms have been associated with the development of the more prevalent, sporadic form of AD, including obesity (*10*), increased plasma cholesterol and type 2 diabetes mellitus (*11, 12*).

The human microbiome has been described to have a distinct, complex role in the pathophysiology of multiple diseases (*13*). Inter-individual variation (*14*) as well as age-dependent changes (*15*) in the human gut microbiome have been described, with Firmicutes and Bacteroidetes identified as the predominant phyla while gut microbial involvement has been suggested in neurodevelopmental (*16, 17*), autoimmune (*18*), movement (*19, 20*) and neurodegenerative (*20, 21*) disorders. Recent studies have proven gut microbial involvement in the development of AD (*22*). Altering gut microbe composition in Alzheimer’s mice via long-term antibiotic administration resulted in decreased cerebral Aβ loading (*23*) while removal and recolonization of gut microbiota resulted in decreased and increased amyloid loading respectively (*24*). Furthermore, the composition of bacterial populations in fecal samples of AD patients is markedly different from healthy individuals in the abundance of certain phyla and genera, with AD patients observed to have increased Bacteroidetes and decreased Firmicutes and *Bifidobacterium* levels (*25*). Hence the involvement of intestinal microbiota in the development and progression of AD has been established, although its exact role has not yet been delineated. FMT involves the transfer of fecal material from a healthy donor to a recipient with the aim of normalizing composition and therefore function of intestinal microbial populations (*26*). FMT has been successfully used to treat recurrent *Clostridium difficile* infections (*27*) with expansions of its application into the treatment of other gastrointestinal disorders (*28*). The efficacy of FMT has also been described in reducing intestinal dysbiosis in Parkinson’s mice (*9*) and alleviating psychiatric disorders (*29*), although its effects in AD has not as of yet been described.

To understand the impact of FMT in a murine model of AD, we performed the procedure via oral gavage in twenty-three 5xFAD (30-32 weeks) recipient mice using fecal matter from healthy B6SJL wildtype donor mice (10-12 and 30-32 week-old) over a period of seven days. Eight of the old recipient mice received oral gavage of fecal slurry obtained from healthy, wild-type donor mice of the same age (Old Tg-fed old; Old Tg-FO) and eight received oral gavage of fecal slurry from healthy, younger wild-type donors (Old Tg-fed young; Old Tg-FY). The control 5xFAD recipient mice (30-32 weeks old; n = 7) were gavaged with normal saline (Old Tg-Control). Following an incubation period of 21 days, mice were subjected to behavioral testing to study the effects of FMT on cognition, after which cerebral Aβ levels were evaluated histologically. Behavioral studies were also performed on healthy wildtype mice who did not exhibit AD phenotype and served as positive controls for comparison against normal behavior (Old-WT). Cognitive ability was assessed as the degree of anxiety, recognition memory and spatial memory. Emotional symptoms such as anxiety arise from Aβ and tau accumulation in the amygdala (*30*) and can be assessed by means of the elevated plus maze (EPM) (*31*) while dorsal hippocampal involvement in recognition and spatial memory formation and hippocampal degeneration during the course of AD are evaluated by the novel object recognition (NOR) (*32*). Synaptic degeneration of the CA1 field in the hippocampus, innervated by pyramidal neurons from the entorhinal cortex, results in spatial memory decline in AD mice (*33, 34*) which is also apparent in hippocampal atrophy and neuron loss seen in AD patients (*35*) and is evaluated in the form of a forced alternation Y-maze (FAY) (*31*). Following treatment, Old Tg-FY mice were found to have overall improved cognition and Old Tg-FO displayed some cognitive improvements. The increasing trend in cognitive improvement observed in the Old groups correlate with decreased Aβ loading.

The elevated plus maze analysis included measuring the percentage of time spent in the closed arms and the ratio of closed to open arm entries. Mice in the Old Tg-FO group spent, on average ~5% more time in the closed arms (n=8, p=0.687; Student’s t-test) whereas Old Tg-FY mice spent nearly twice as much time in the closed arms (n=8, p=0.056; Student’s t-test) when compared to Old Tg-Control (n=7) animals. Although the time spent by the Old Tg-FY group was not statistically significant it was similar to the time spent by WT mice (n=4), demonstrating a recovery of anxiety that is lost during the disease process (Fig 1a) (*36*). Old Tg-FO mice also demonstrated change (p=0.911) in the ratio of closed arm to open arm entries while Old Tg-FY showed a small increase in this ratio (p=0.477), with levels somewhat identical to age-matched WT littermates (n= 4; Fig 1b) (*36*).

Examining performance in the Novel Object Recognition test, Old Tg-FO mice demonstrated more than a two-fold increase in the Discrimination Index (DI) (p=0.065), indicating a greater preference for the Novel Object when compared to Old Tg-Control mice. Old Tg-FY demonstrated a significantly greater, approximately five-fold increase in the DI (p=0.02), indicating a much higher preference for the Novel Object compared to control (Fig 1c) (*36*).

The FAY results were analyzed in the form of time spent in the Novel Arm and ratio of Novel to Other arm entries. Old Tg-FO animals showed a significant, near two-fold increase (p=0.046) while Old Tg-FY demonstrated a significant two-fold increase (p=0.030) in time spent in the Novel Arm (Fig 2d). Compared to the Old-WT mice who were used as a standard, the Old treated groups spent even more time exploring the Novel Arm. When comparing the ratio of Novel:Other arm entries, Old Tg-FO mice also displayed an approximately two-fold, statistically significant increase (p=0.012) in preference for the Novel arm while Old Tg-FY mice demonstrated a nearly three-fold, statistically significant increase (p=0.012) in preference (Fig 2e) (*36*). Compared to the Old-WT mice, both groups of Old-treated animals showed a higher sense of exploratory behavior, demonstrated by a greater ratio of Novel:Other arm entries (Fig 1e) (*36*).

Aβ plaque loading was evaluated using Thioflavin S staining and analyzed as the percentage area of cortical amyloid plaque and quantified as the number of cortical plaques. Old Tg-FO had a decrease (p=0.098) in the area of amyloid plaque present in the cortex while Old Tg-FY had a remarkable nearly two-fold decrease (p=0.002) when compared to Old Tg-Controls (Fig 1f). Both Old Tg-FO and Old Tg-FY demonstrated a decrease in the number of quantifiable plaques within the cortex, with Old Tg-FO having a near 2.5-fold decrease (p=0.060) and Old Tg-FY having more than a 2.5-fold decrease (p=0.045) (Fig 1g). As both plaque size and number of plaques decreased notably, overall amyloid plaque loading can be interpreted as having decreased (Fig 1i) (*36*).

The decreasing trend of Aβ plaque load across Old Tg-FO and Old-Td-FY is inversely correlated with the trend of improved cognition, which was measured as a compounded Cognition Score (CS) of the behavioral study results, indicating the importance of donor age in decreasing amyloid pathology and improving cognition (Fig 1h) (*36*)

During the EPM, mice were placed in the middle platform and allowed to explore for 5 minutes. The percentage of time spent in the closed arm and ratio of closed arm entry to open arm entry was measured as indicators of anxiety (Fig 1a). Old Tg-FO mice showed no discernible improvement in anxiety levels, indicating little to no improvement of amygdala deterioration. Old Tg-FY mice showed a trend of improvement, indicating the possibility of reduced anxiety levels. The amygdala interacts with the hippocampus and hypothalamus to influence the hypothalamic-pituitary-adrenal axis which is involved in the regulation of the stress hormones such as cortisol. A reduction of dysbiosis of intestinal microbiota could result in restoration of permeability of the intestinal epithelium and possibly the normalization of cortisol levels via the gut-brain axis, causing a decrease of anxiety levels, especially in Old Tg-FY mice. Furthermore, FMT may allow for increased clearance or renewal of clearance mechanisms of Aβ from the amygdala which could result in reduced anxiety. The overall results of the EPM show a trend towards normalization of anxiety to levels similar to that observed in age-matched wildtype mice.

The NOR involves exposure to an object (Familiar Object) for 5 minutes in Phase 1, followed by an interval and subsequently, exposure to the Familiar Object and a Novel Object for 5 minutes. To understand object preference in the mice, we calculated the DI which assigns a numerical value indicating object preference. The more negative the number, the lesser the preference for the Novel Object. Old Tg-FO and Old Tg-FY groups both demonstrated increases in the DI, indicating improved recognition memory sensitivity (*37*) in the treated groups compared to controls. Our results indicate the restoration of hippocampal function to a certain extent in treated groups, possibly related to increased hippocampal Aβ clearance. Treated groups demonstrate a reversal in recognition memory deficits to levels comparable to wildtype littermates in the NOR.

The procedure for the FAY was similar to the NOR, involving an initial phase where mice were given 5 minutes to explore the 3-arm maze with a single arm blocked from entry. In Phase 2 mice had access to all 3 arms. Old Tg-FO and Old Tg-FY showed remarkable improvement in spatial memory following FMT, with increases in both the percentage of time spent in the Novel Arm and the ratio of Novel arm entries to other arm entries (Fg 1d-e). The FAY results demonstrate that FMT is particularly efficient in improving spatial memory, indicating its potency in restoring hippocampal function. Similar to the EPM and NOR results, the FAY outcomes demonstrate that FMT may be facilitating Aβ clearance, allowing for improved spatial memory.

Decrease in Aβ levels was investigated quantitatively using Thioflavin S staining. Thioflavin S binds to amyloid and is a widely used method for cortical Aβ staining (*38*). The percentage area of cortical plaques and the number of plaques were calculated to understand Aβ plaque loading in treated and control mice. The percentage area can be interpreted as the size of cortical Aβ plaques and the decrease in area indicates shrinkage of cortical amyloid plaques. This could be attributed to increased clearance and reduced deposition of Aβ, possibly by FMT-induced renewal of immune-mediated clearance mechanisms or inflammatory regulation (*39*). To reflect the correlation of Aβ plaque loading and cognitive performance, a Cognition Score (CS) was calculated for each animal in each group as a compounded score of the behavioral studies results and plotted against the mean percentage area of amyloid plaque, or amyloid plaque loading (AP). The resulting correlation showed both Old Tg-FO and Old TG-FY having higher cognition with lesser Aβ plaque burden compared to the Old Tg-Control. Interestingly, Old Tg-FY had a much higher CS and lower AP compared to Od Tg-FO, indicating that donor age influences FMT outcomes.

Restoring gut microbial compositions and reducing dysbiosis is a key function of FMT. Young wild-type (10-12 weeks) donor mice would have distinct microbial compositions compared to that of Old wild-type mice (30-32 weeks) (*40*) and recipient mice gut microbial composition would be restored to that similar of donor mice. Microbiota from aged mice contributes to low-grade inflammation (*41*), hence normalizing gut microbe compositions to a younger population would result in the reduction of intestinal and systemic inflammation. Consequently, neuroinflammation-associated deterioration, such as the loss of blood-brain-barrier integrity (*42*), could be reversed. Dysfunctional metabolite levels in AD (*43*) could be repaired by FMT, culminating in the normalization of metabolite levels affected by gut microbial function and composition and subsequent reduction of pathology. The gut-brain axis (GBA) comprising of bidirectional communication between the enteric nervous system (ENS) and the central nervous system (CNS) such as neurotransmitter expression, anxiety and stress levels, and cognition is also affected by intestinal microbes (*44*). With the marked decrease in amyloid loading of the treated mice, we conclude that FMT was able to increase clearance of cortical Aβ, possibly via immune or inflammatory regulation, allowing for improved cognition.

## Author contributions

Ms Elangovan and Dr Holsinger were involved in conceptualization, data curation, formal analysis, investigation, methodology, project administration, validation and visualization of the project. Dr Holsinger supplied resources, software, supervision and writing – review and editing. Ms Elangovan prepared the original draft. Professor Borody assisted in interpretation of the data and provided critical revision of the manuscript.

## Data and materials availability

Analyzed data is available in the main text or the supplementary materials. All data is available as on the Research Data Store (RDS) of the University of Sydney.

## Supplementary Materials

### Materials and Methods

**Fig. S1.**
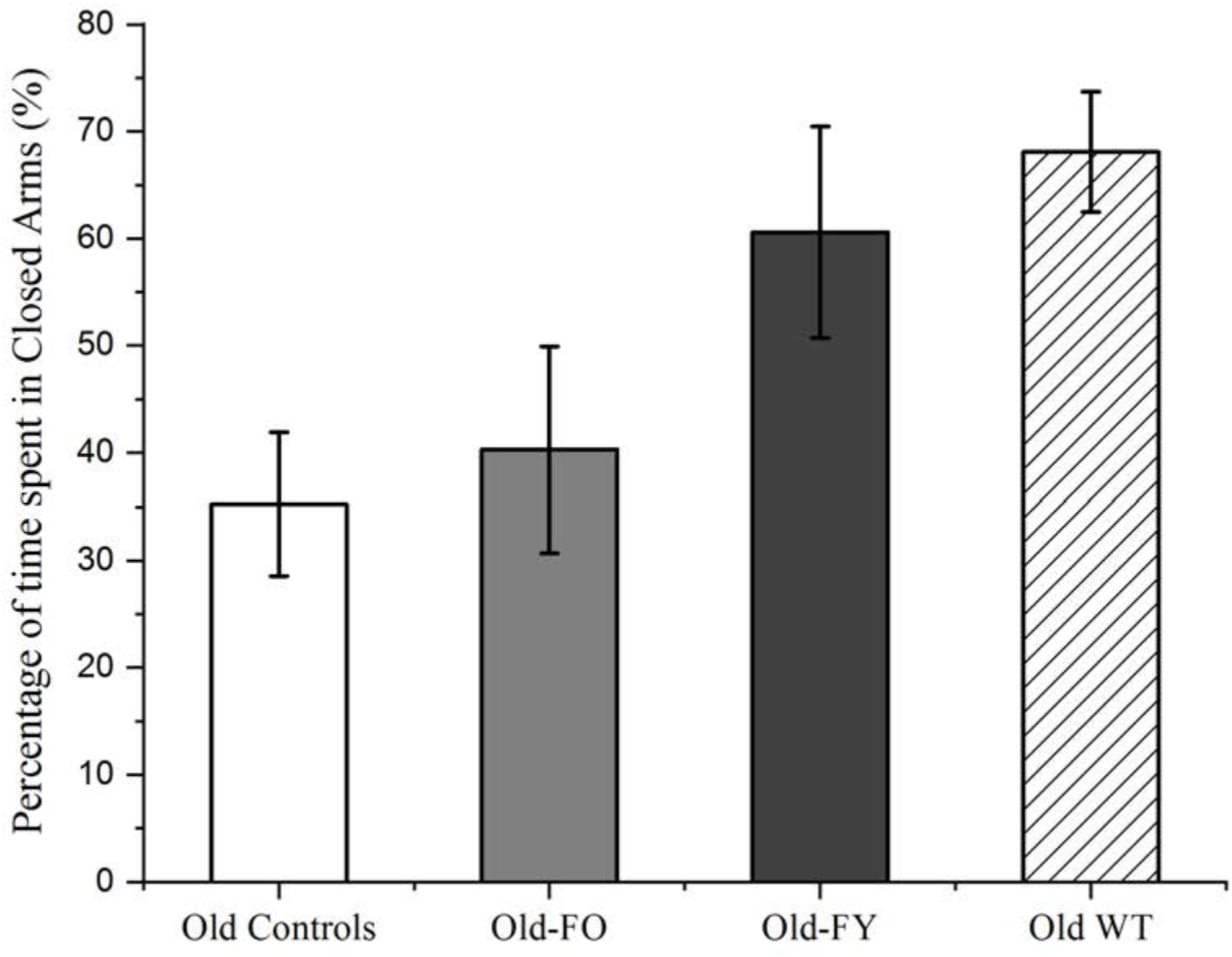
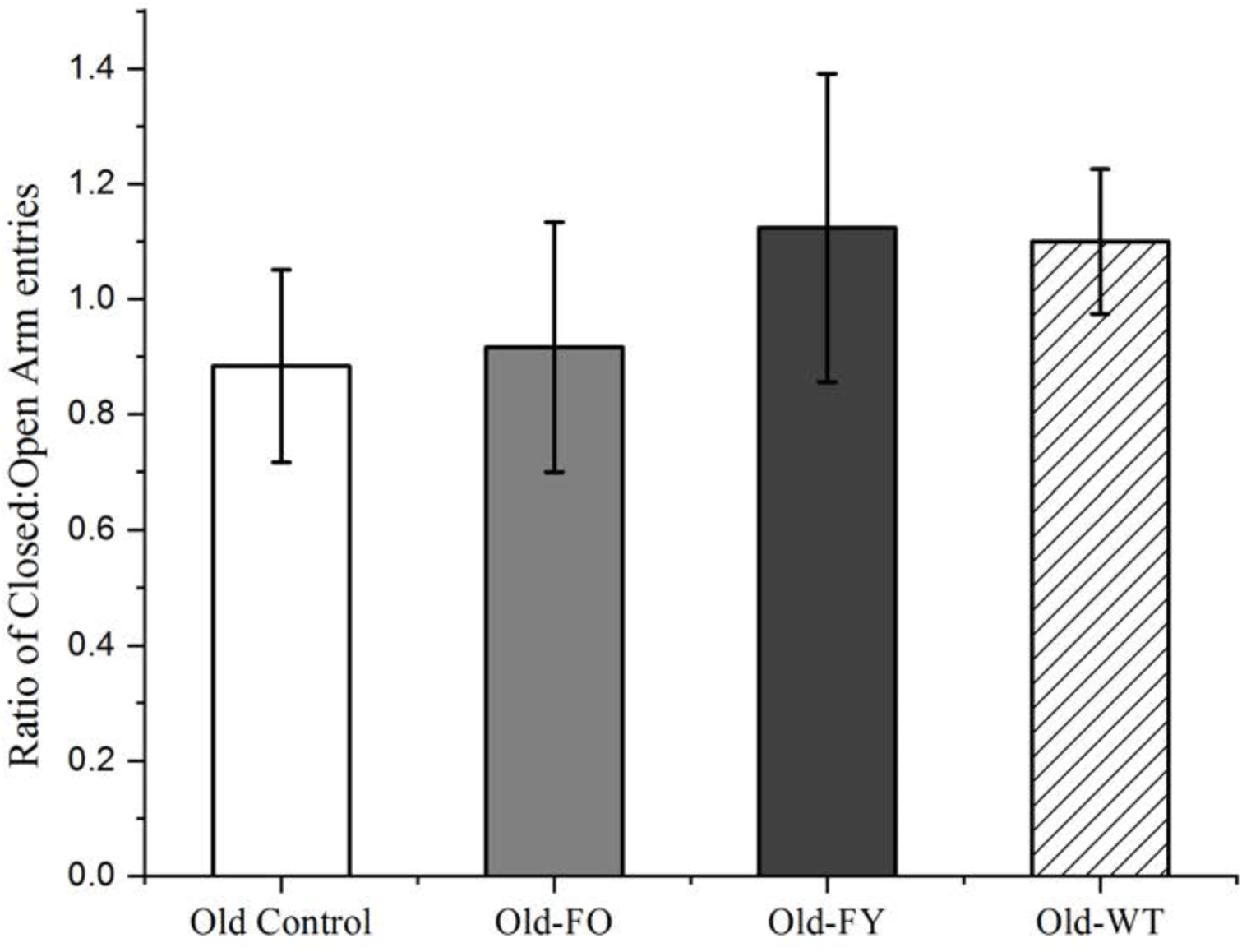
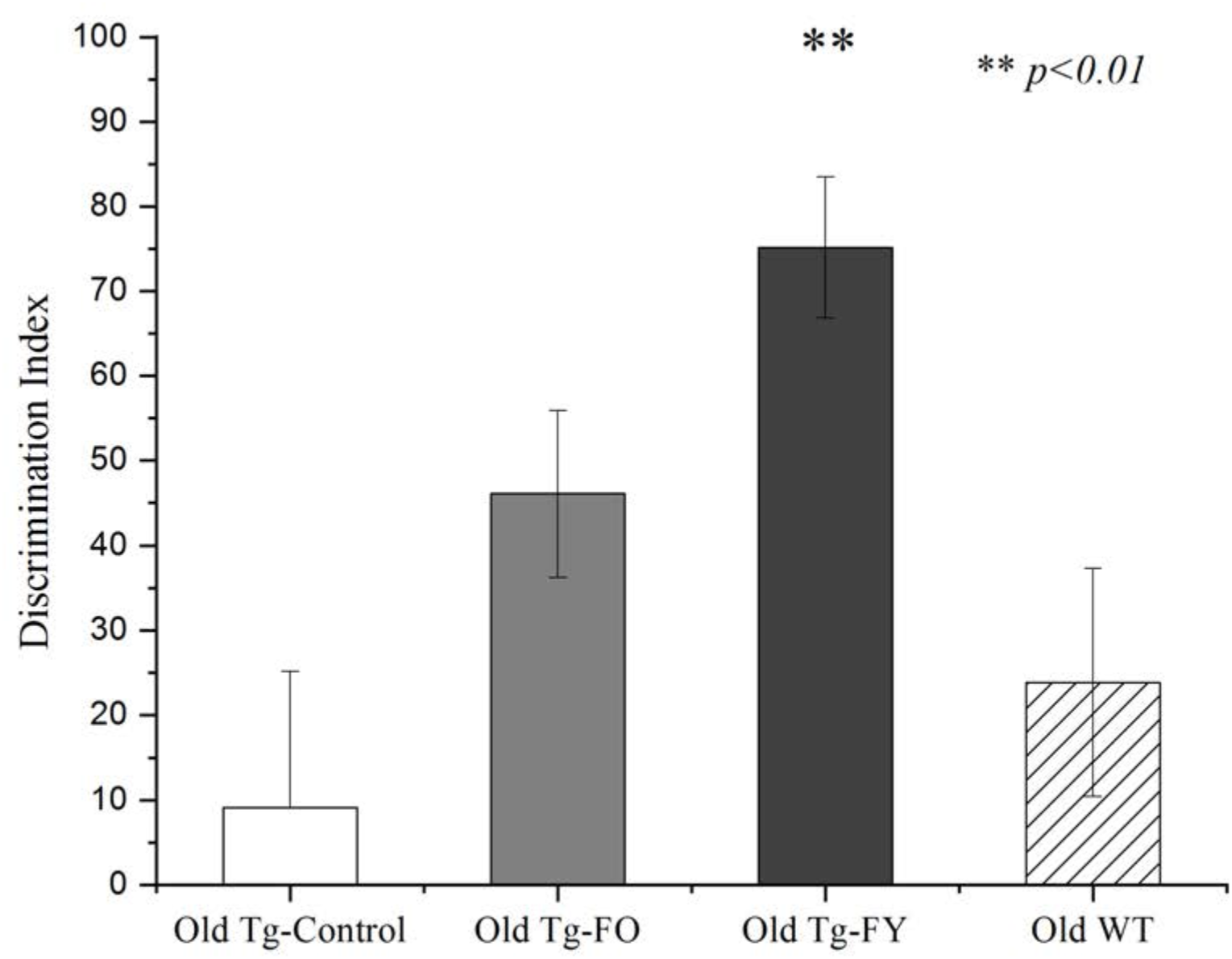
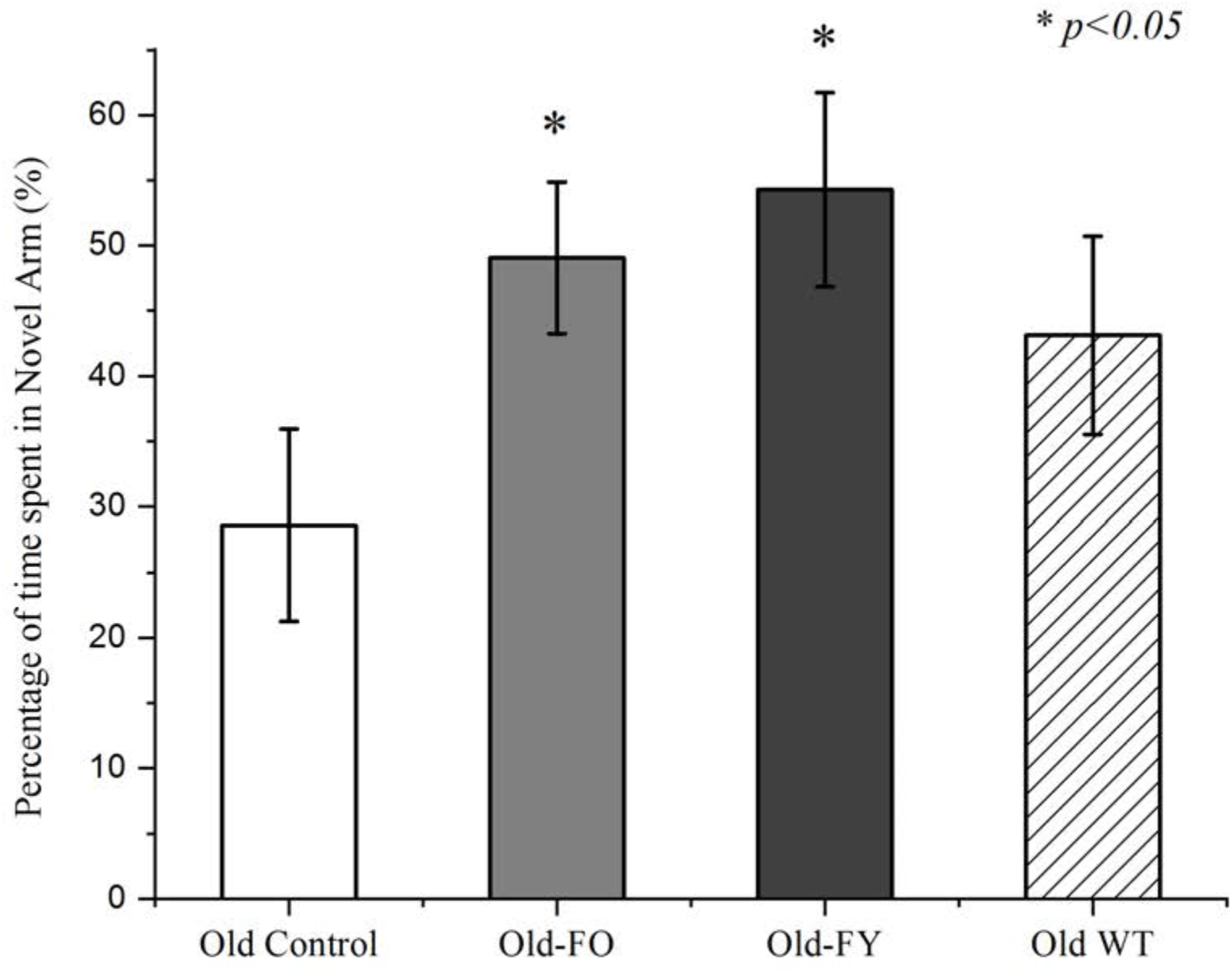
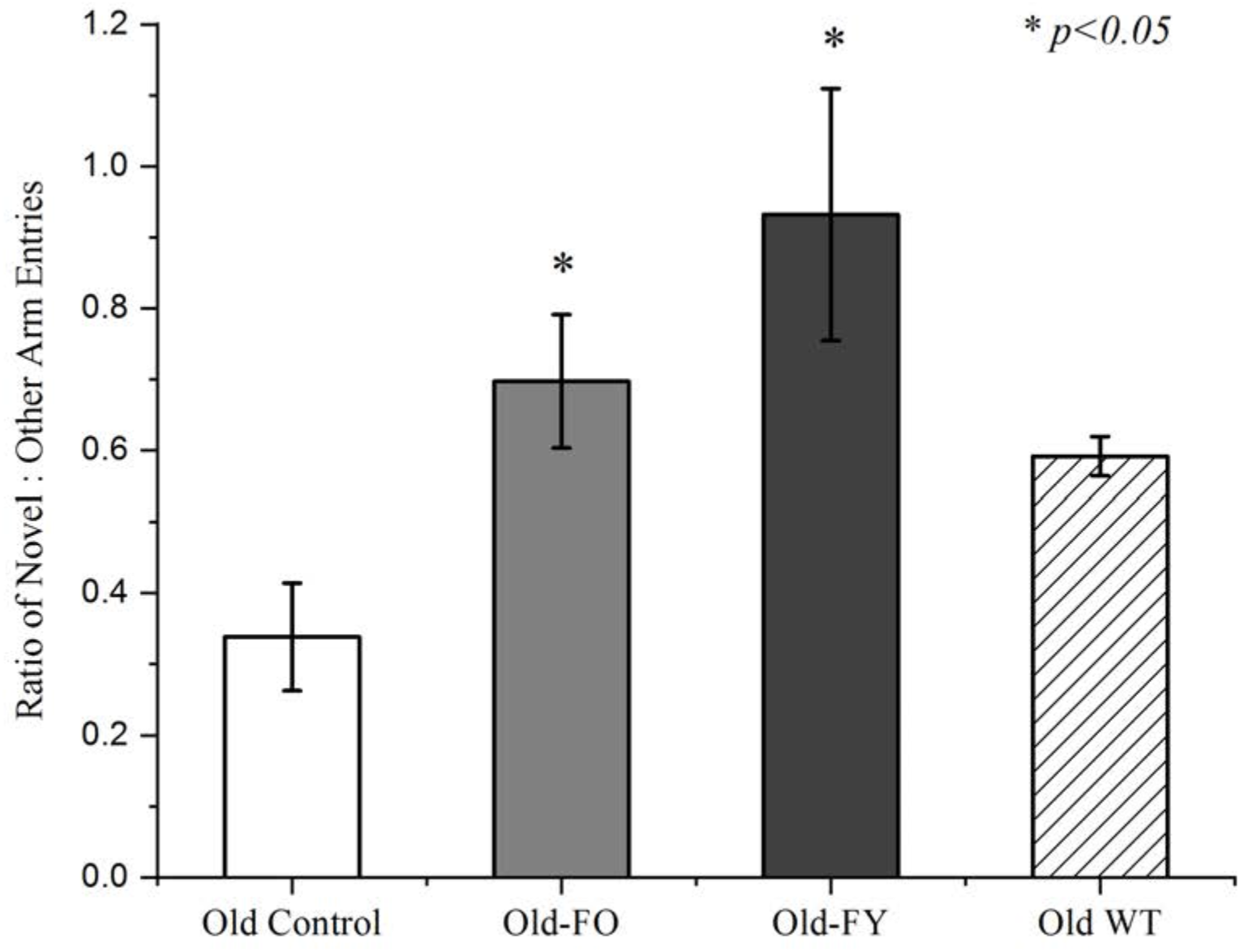
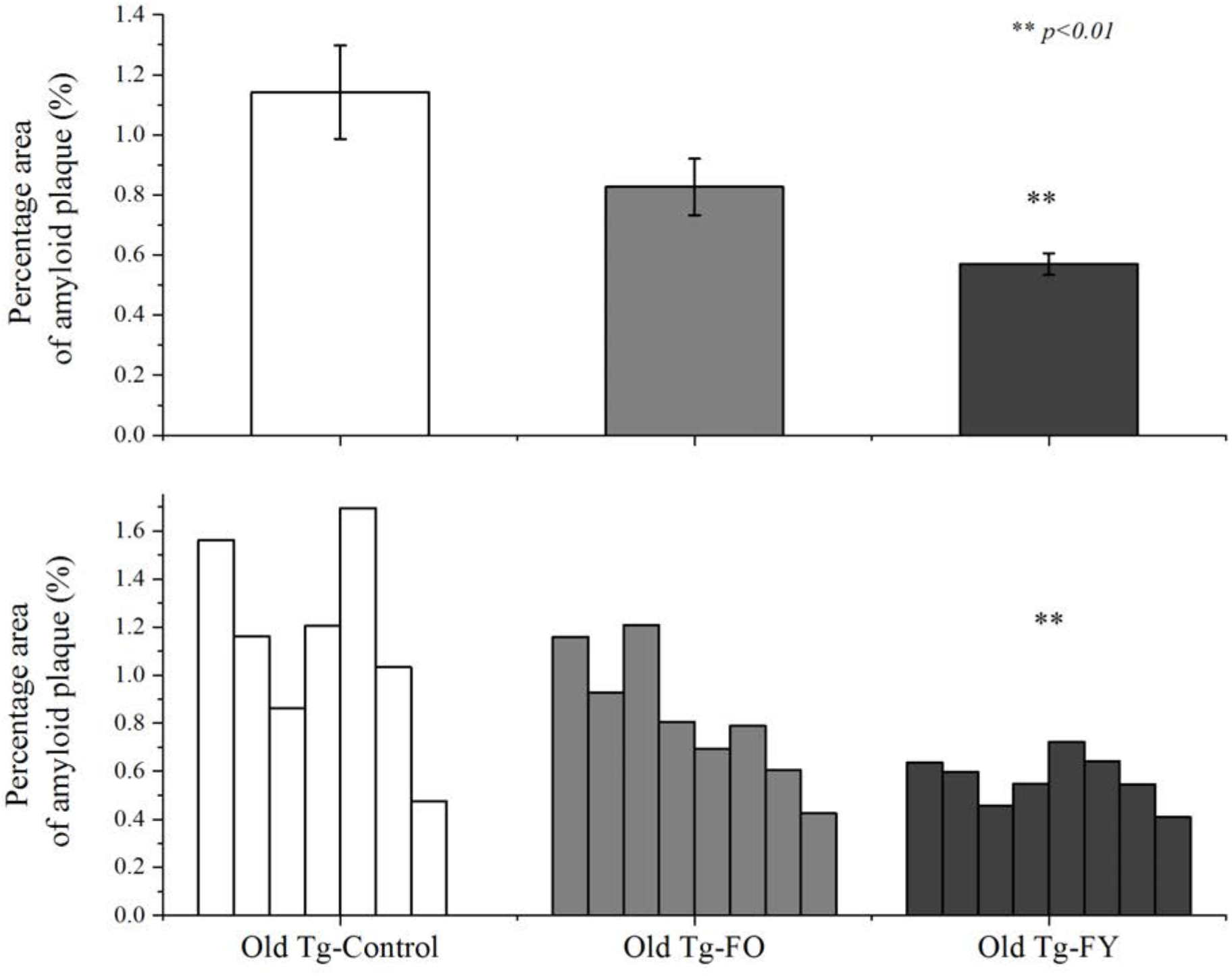
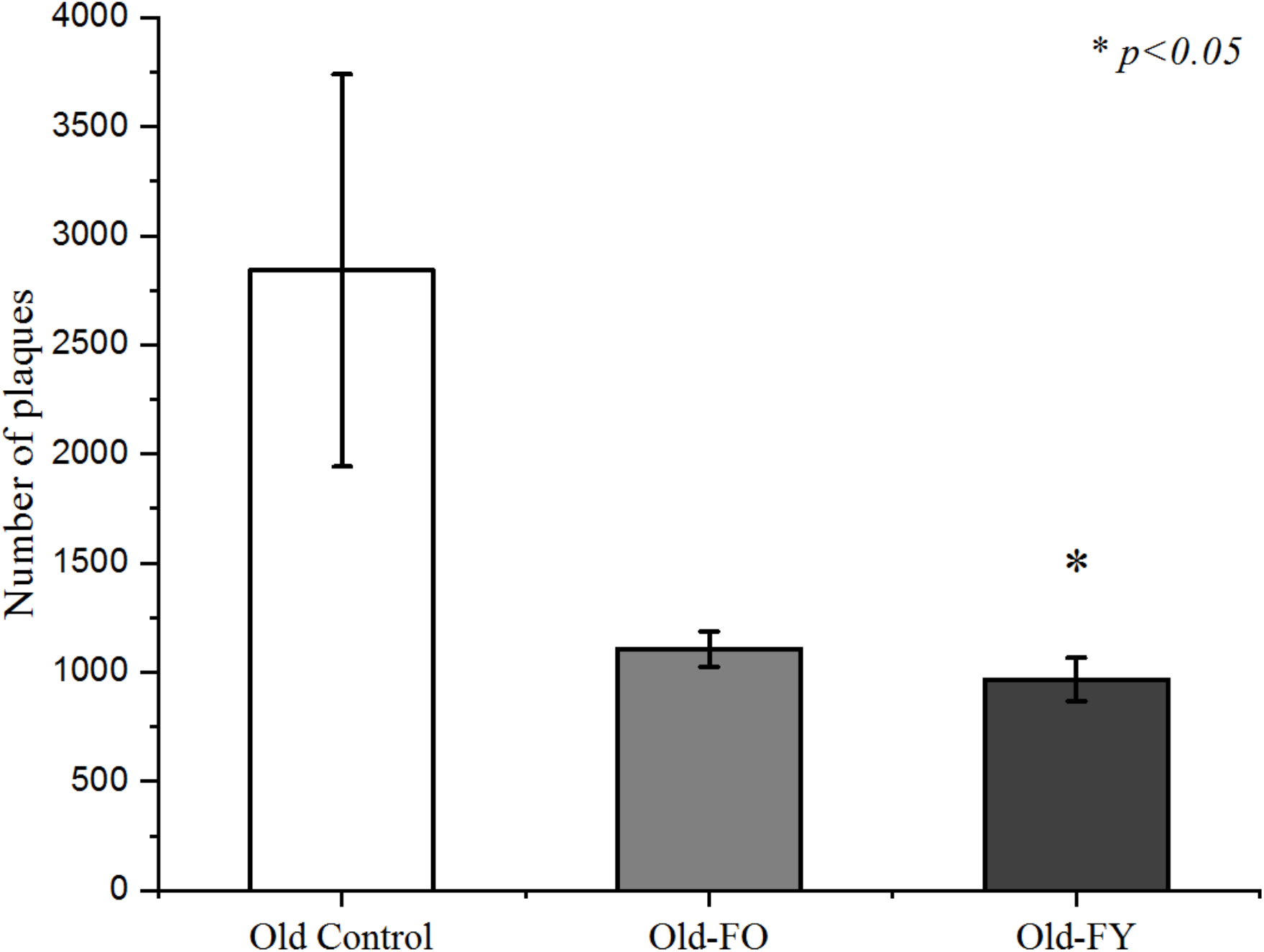
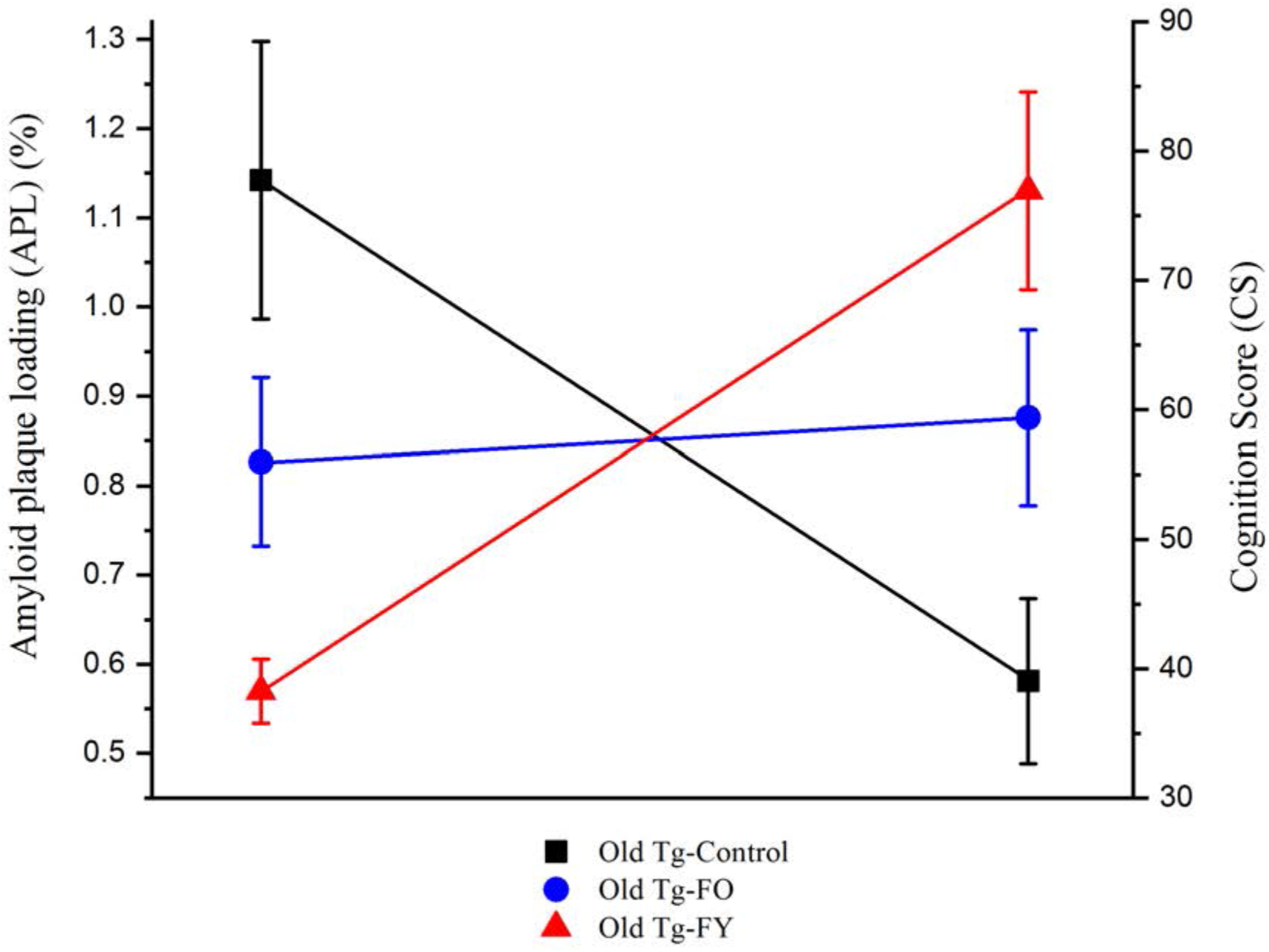
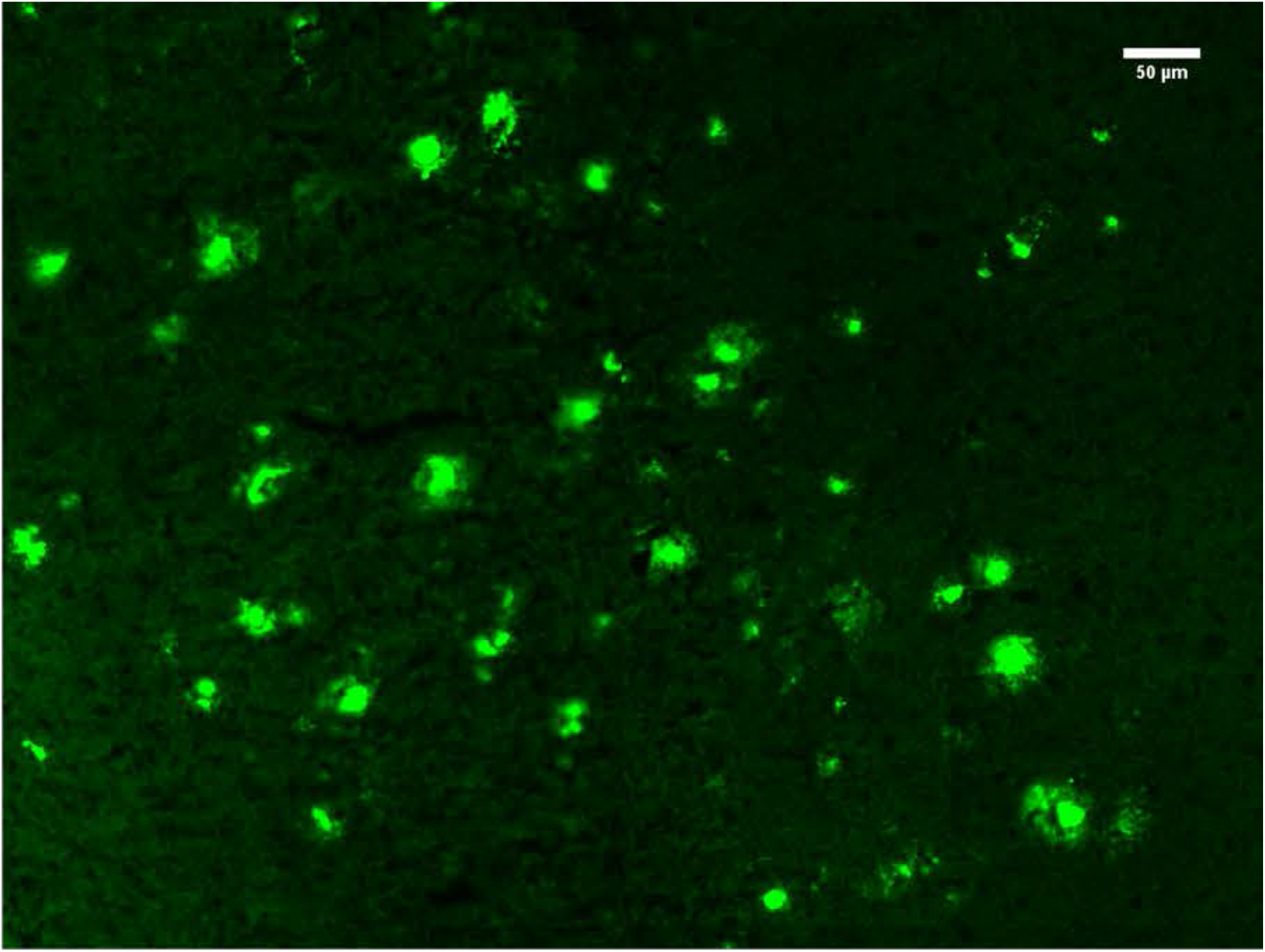
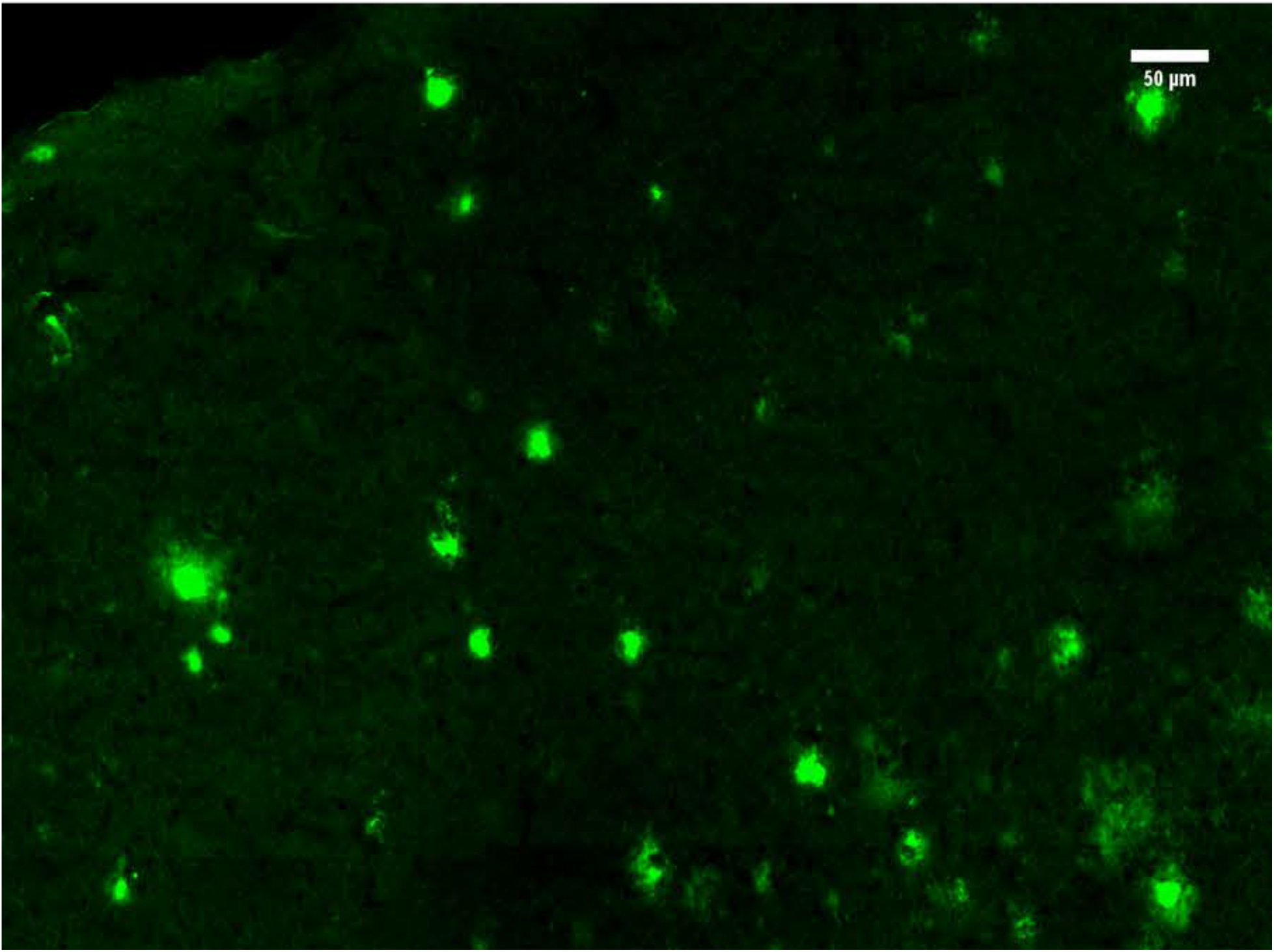
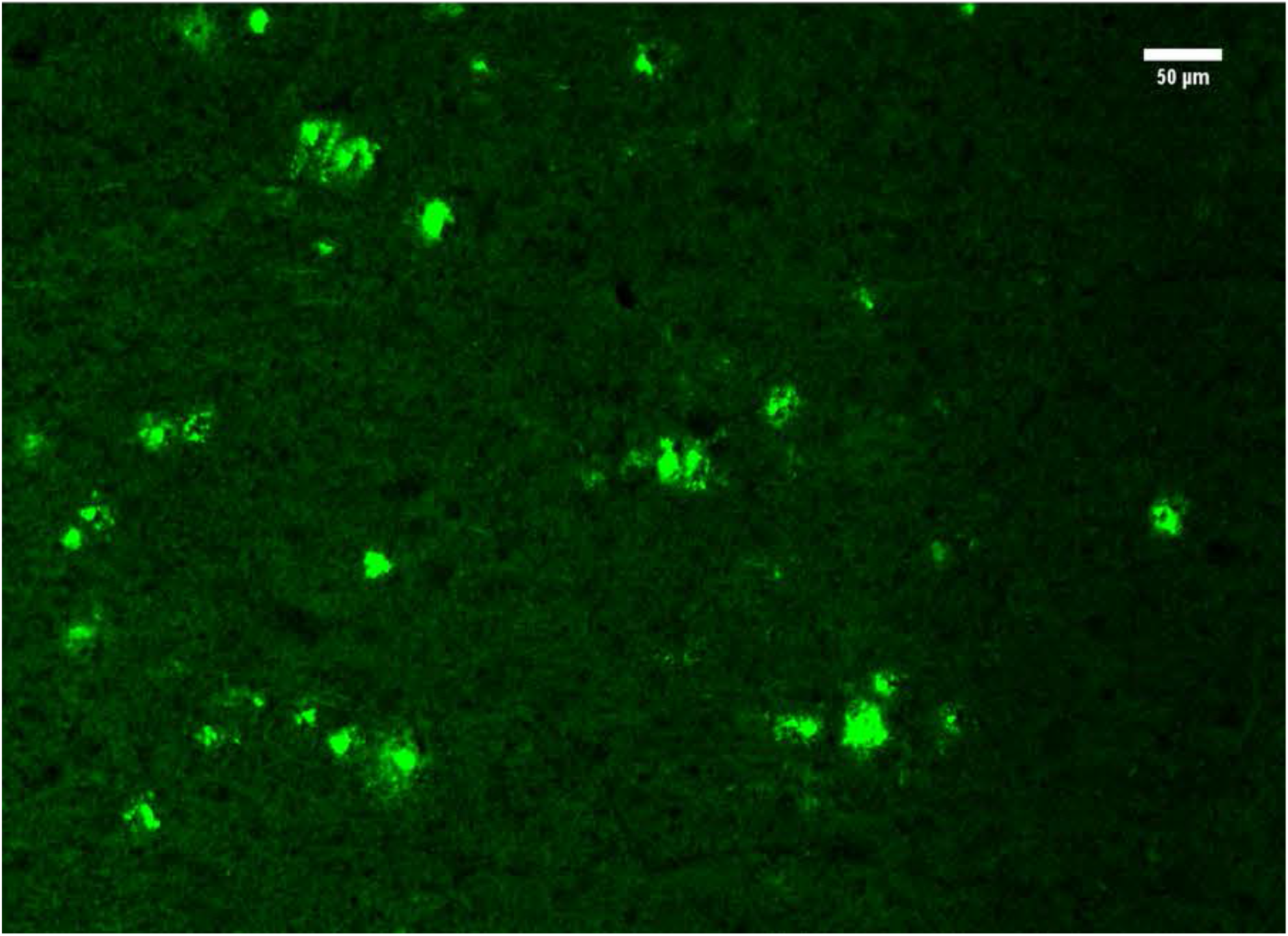
(A) Percentage of time spent in closed arms. Data is presented as mean ± SEM. WT values are presented for comparison only. (B) Ratio of Closed:Open arm entries. Data is presented as mean ± SEM. WT values are presented for comparison only. (C) Discrimination Index. Data is presented as mean ± SEM. WT values are presented for comparison only. **p<0.01 (D) Percentage of time spent in Novel Arm of Y-maze. Data is presented as mean ± SEM. WT values are presented for comparison only. * p<0.05 (E) Ratio of Novel:Other Arm entries in Y-maze. Data is presented as mean ± SEM. WT values are presented for comparison only. * p<0.05 (F) Percentage area of cortical amyloid plaque. Data is presented as mean ± SEM. **p<0.01 (G) Number of quantifiable cortical plaques. Data is presented as mean ± SEM. *p<0.05 (H) Cognition Score against Amyloid Plaque Loading (I) Thioflavin S stain of amyloid plaques in the cortex of an Old Tg-Control mouse (J) Thioflavin S stain of amyloid plaques in the cortex of an Old Tg-FO mouse (K) Thioflavin S stain of amyloid plaques in the cortex of an Old Tg-FY mouse

Ethics approval was obtained from the University of Sydney Animal Ethics Committee (AEC Number: 2017/1285) and all procedures were in accordance with institutional guidelines. Heterozygous male and female 5xFAD mice overexpressing three human APP (Swedish, Florida and London) FAD mutations as well as two human Presenilin 1 mutations (*37*) were bred under standard laboratory conditions at the Bosch Laboratory Animal Services Facility and were maintained on a 12:12hr light/dark cycle with food and water provided *ad libitum*. Wild type littermates served as fecal donors.

#### a. Fecal microbiota transplant

Fresh fecal matter was collected directly from healthy wildtype donors aged 10-12 (Old Tg-FY) and 30-32 (Old Tg-FO) weeks. Fecal matter was homogenized using a micro-centrifuge pestle into a 50mg/ml fecal slurry using 0.9% sterile saline solution. The slurry was then micro-centrifuged three times with collection of the supernatant and resuspension of the pellet each time prior to centrifuge. The pellet was discarded while the supernatant was retained and used for oral gavage. Each treated mouse received 200μL of supernatant via oral gavage using a 22G oral gavage needle daily for 7 days with control animals receiving 200μl of sterile saline. Mice were monitored for general health and weight during the gavage procedure. Following oral gavage, all mice underwent a 3-week incubation period with food and water available *ad libitum*.

#### b. Behavioral studies

All testing was video-recorded. Mazes were wiped with F10SC disinfectant after each trial to remove odor cues. Mice were handled for 3 days prior to testing to allow for acclimatization.

##### i. Elevated Plus Maze (EPM)

The elevated plus maze consists of 4 arms of equal length joined by a central platform with 2 opposing arms enclosed by walls with the maze elevated 1m off the floor. Mice were introduced to the maze for the first time during testing and were given 5 minutes each to explore the maze. An arm entry was recorded when 80% of the mouse’s body was in the arm. Stretch-attend posture was not included as an arm entry. Time spent in the central platform was not recorded. Arm entries were measured and analyzed as a ratio of entries into closed arms to entries into open arms. The duration of time spent in the open and closed arm each was measured in seconds and analyzed as a percentage of time spent in closed arms.

##### ii. Novel Object Recognition (NOR)

The NOR test consisted of 2 phases separated by a 4-hour interval and was conducted in a box measuring 1m by 1m. Mice were acclimatized to the empty box for 5 minutes each, 24 hours prior to testing. During Phase 1 of testing, mice were given 5 minutes to explore the box with an object (Familiar Object) affixed to the floor. Interaction with the object was defined as sniffing, pawing and interaction with the object and was counted when the animals directed their nose within 1cm of the object. Mice were then removed and the box and Familiar Object were disinfected using F10SC disinfectant. In Phase 2, 4 hours later, mice are reintroduced to the box, with a second object (Novel Object) affixed to the floor in a new position in addition to the retention of the Familiar Object in the original position. The time spent interacting with the Novel Object and Familiar Object in Phase 2 was measured separately in seconds. Novel object interaction was defined as a Discrimination Index calculated as the difference between time spent exploring the Novel and Familiar Object as a percentage of the total time spent exploring both objects.

##### iii. Forced alternation Y-maze

The forced alternation Y-maze consisted of 2 phases separated by a 4-hour interval and was conducted on a maze comprised of 3 enclosed arms of equal length joined by a central platform. Mice were placed in the central arm at the start of Phase 1 and given 5 minutes to explore the maze with entry to one arm blocked. Visual cues were placed on the walls above the closed arm. Mice were then returned to their cage and the Y-maze was disinfected. Following the inter-trial interval of 4 hours, mice were returned to the maze for Phase 2, where each mouse was allowed 5 minutes of exploration with all 3 arms open, with the newly opened arm referred to as the Novel Arm. The same visual cues were in place in Phase 2. The duration of time spent in the maze was analyzed for Phase 2 only and the time spent in each arm in seconds as well the number of arm entries were recorded. An arm entry was counted when 80% of the mouse’s body entered the arm. Stretch-attend posture was not included as an arm entry. Time spent in the central platform was not counted. Novel arm entries were analyzed as a ratio of novel arm entries to other arm entries and time spent in the Novel Arm during Phase 2 was measured in seconds and analyzed as a percentage of time spent in all arms.

#### c. Cognition Score

A Cognition Score (CS) was calculated by assigning indexes measured out of 100 to each of the 5 parameters measured for each group of mice and averaging the score. The resultant CS is an index out of 100 indicating overall cognitive performance.

#### d. Thioflavin S staining and immunofluorescence detection

Mice were deeply anesthetized using isoflurane and decapitated. The brains were extracted, snap frozen in liquid nitrogen-cooled isopentane and stored at −80°C. Six, 16μm coronal slices (from Bregma Zero to Bregma −1.28mm) for each mouse were obtained via cryosectioning and were stored on glass slides at −20°C. Thioflavin S (0.015%, Sigma-Aldrich Inc, Seven Hills) was prepared as a solution in 50% ethanol. Slides were thawed to room temperature for 10 minutes followed by 10 minutes in an acetone wash at −20°C. The slides were then washed in phosphate-buffered saline for 5 minutes and Thioflavin S solution was placed directly on each section and slides were incubated for 10 minutes in a dark humidity chamber. Slides were then washed twice in 80% ethanol for 2 minutes each and once in 95% ethanol for 1 minute followed by 3 washes in distilled water for 1 minute each. Sections were sealed under coverslips with Fluoromount mounting media (Sigma-Aldrich Inc.). Images were obtained at 20x using the Zeiss Axio Scan.Z1 slidescanner system. Fiji image processing software (NIH, ImageJ) was used to analyze Thioflavin S staining. The Shapiro-Wilk test of normality and a Student’s t-test for analysis of images was performed using the SPSS Statistics Package (version 24; IBM).

